# Pitfalls in understanding PhIP-Seq data: technical variability, replicability, and best practices for interpretation

**DOI:** 10.64898/2026.09.15.751724

**Authors:** Lovro Trgovec-Greif, Thomas Vogl

## Abstract

Phage Display Immunoprecipitation sequencing (PhIP-Seq) is a high throughput method allowing to measure antibody binding against hundreds of thousands of potential antigens. Typically, blood samples of hundreds to thousands of individuals are measured in 96-well plates, necessitating distribution of samples on multiple plates and processing in batches. To correctly interpret PhIP-Seq results, it is crucial to understand its nature and technical limitations.

Here, we analyzed 195 technical replicates of controls (“anchor samples”) from 49 plates measured in 13 immunoprecipitation runs to gauge technical variability and replicability. While there were few false-positive enriched peptides, we observed a substantial fraction of false negatives. The total number of enriched peptides can be affected by batch effects. Additionally, the type of biological material used (serum or plasma) influences enrichment profiles with an antigen library containing bacterial proteins due to the interaction of phages with plasma proteins. Finally, replicating measurements in different labs produces comparable enrichment profiles, but slight differences in a subset of peptides are noticeable.

These results suggest best practices for PhIP-Seq experiments. Most importantly, due to batch effects, samples of cases and controls should be evenly distributed between 96-well plates to avoid confounding with biological effects. Also, for highly complex libraries, the total number of enriched peptides can be a technical artifact and potentially not a feature useful in comparisons.

## Introduction

Phage display immunoprecipitation and sequencing (PhIP-Seq) is a technology for high throughput profiling of the antibody binding capacity of complex samples (for example, human serum). Current technology allows for probing the IgG (most common), IgA [1] and IgE [2, 3] antibody repertoires. Initially developed [4] to study autoimmunity (for example, [5–8]), its use has been extended to study antibody repertoires against viruses [9, 10], the microbiome [11], toxins, allergens [2, 12], and others (for a comprehensive overview see ref. [13]). PhIP-Seq is based on encoding peptide antigens as synthetic DNA, their display on phages, incubation with the sample containing antibodies, precipitation of the phages bound by antibodies and subsequent sequencing to detect enrichment. PhIP-Seq can be scaled on liquid handling robots, and typically, blood samples of hundreds to thousands of individuals are measured in 96 well plates, therefore necessitating distribution of samples between multiple plates and processing in batches.

While powerful in its ability to detect antibody binding against peptides of interest, PhIP-Seq is not a widely used technique such as ELISA, and so far, no systematic exploration of its limits and peculiarities has been made available. Knowledge of the extent of common errors such as false positive and false negative binding events (“hits”) as well as replicability and the extent of batch effects [7] has implications for downstream data analysis and required cohort size to answer a question of interest. On top of the intrinsic properties of the method influencing the errors and variability, every lab has its own implementation of the assay potentially further impacting the results and complicating comparisons. Some effects will remain lab specific unless the method gains widespread popularity which would lead to more standardization in terms of procedures and the analysis software [14] and possibly production of commercial kits.

Our lab routinely conducts PhIP-Seq experiments [15–18] on a liquid handling robot and, we have accumulated data for 195 replicates of control (“anchor”) sample from 49 experimental plates processed in 13 batches spanning around 2 years of measurements. Our experiments probe the influence of various factors such as the laboratory conducting measurements and sample type on the assay results. These data offer an opportunity to understand the intricacies of PhIP-Seq data as well as to learn about various pitfalls in the downstream analysis due to assumptions about the data that may bias the interpretation. Here, we report replicability, stability and validity of the data generated by PhIP-Seq based on these anchor sample replicates. As the anchor sample was measured in multiple runs at different times, we also detected batch effects associated with the experimental 96-well plate and the immunoprecipitation (IP) run. Besides showing the results obtained by repeatedly measuring the anchor sample, we also show that the biological material used for the experiment (serum or plasma) influences the ability to detect antibody binding to peptides that interact with coagulation factors with this specific antigen library. Additionally, we compared the same set of samples measured with the same library in two different labs. While the results from both labs match in general, there is a small number of hits that are absent among the results from one or the other lab.

**Motivation**

Writing of this manuscript was partially inspired by the transcript of a speech about “Cargo Cult Science” given by physicist Richard Feynman [19]. Discussing the importance of scientific integrity, rigor, and honesty, he gives the example of troubleshooting experiments on rats running in a maze. A researcher wanted to train rats to enter a specific door in a maze made of corridors by conditioning them with food. For some reason, the rats could always tell where the food had been the last time, and would enter that door (rather than entering the door the researcher intended to train them). The researcher had to find out what property of the maze itself allowed rats to find the food. Finally, he discovered it was the sound produced while the rats were running that revealed which door the rats had to open. He therefore had to first learn a lot about his experimental setup before he could draw any conclusions about the ability of rats to find food in the maze. Similarly, while working on various projects involving PhIP-Seq, learned a lot about the nature of the data and the inherent limitations. We believe it is useful to let the scientific community know what we learned to allow for modification and better planning of the experiments and for improved interpretation of the data.

## Methods

### Principles of PhIP-Seq

In a PhIP-Seq assay, proteins against which antibody binding is to be tested are first rationally selected and their amino acid sequences are obtained either from public databases such as NCBI RefSeq [20] or from a metagenomic sequencing experiment [11]. In brief, proteins are then computationally cut into smaller peptides (for example, 64 aa) with an overlap (for example spanning the length of a linear epitope with a buffer, e.g., 20 aa). The obtained peptides are computationally reverse translated with codon usage optimized for *Escherichia coli* and the oligonucleotides are synthesized (optionally with an identification barcode [21]). The oligonucleotides are cloned into T7 phages for surface display (Novagen’s T7Select® Phage Display System) and expanded in a laboratory strain of *E. coli*. Phages now express the cloned peptide on their surface and the collection of the phages displaying the selected peptides is called a PhIP-Seq library. Phages are mixed with a serum sample (or another antibody containing sample) to allow the antibodies to bind to the displayed peptides and the unbound phages are subsequently washed away. The barcodes [21] of the remaining phages are sequenced, the reads belonging to individual clones are counted and the data are subjected to statistical analysis to detect enriched phages because unbound phages are not washed away completely. When a phage clone is enriched, it is due to the binding of the antibody to the displayed peptide.

### Our implementation of PhIP-Seq

We have reported our protocol in detail in refs. [11, 17] as well libraries used in refs. [11, 12, 22]. In short, when we conduct a PhIP-Seq experiment, we start by mixing phages and serum to allow for antibody-antigen binding. The mixture is then incubated with magnetic beads coated with protein A/G [23] that bind the IgG antibody’s constant region. By using a liquid handling robot (Tecan Fluent 1080), the beads are washed twice with a buffer solution. The remaining phages are then processed as described in [24] and sequenced on an Illumina machine. The current libraries contain antigens from the microbiome and pathogens [11], allergens and toxins [12] and coronaviruses [22]. The size of the combined library is 357 192 peptides.

### Control peptides

In every library, a number of control peptides is included [11, 12, 22]. Positive controls are those against which the antibodies are expected to be present in almost all sera. They represent peptides from common pathogens such as the Epstein-Barr virus, Rhinovirus, Enterovirus or Human adenovirus. Negative controls are peptides against which it is very unlikely to find antibodies. For examples peptides from the Ebola virus or ribosomal and histone proteins. Random controls are the peptides with a random amino acid sequence. No antibody binding to the random controls is expected.

### Terminology

PhIP-Seq is gaining popularity, but is not a routine method and the readership might not be familiar with the terminology. The most important terms and their explanations will be given to make the reading easier. The PhIP-Seq *library* is a collection of all peptides that are expressed on the surface of the phage. A *peptide* is one library component. However, we also use the term library for a collection of the phages expressing library peptides e.g. the physical suspension of phages. The context defines what we are referring to. *Input* is also occasionally used to refer to the library, but rather the physical material of phages used in the experiments. *Input counts* are the number of reads that map to a specific library member after sequencing the library itself. The physical library consists of phage *clones*. Clones are the phages expressing identical peptides and having the same barcode. The number of different clones in a library is identical to the number of uniquely barcoded peptides in the library. In PhIP-Seq, serum antibodies are incubated with the phages to allow for *binding* of the antibodies to the expressed peptides. The phages (clones) expressing a peptide bound by antibodies are *enriched* after immunoprecipitation. As clones are defined by the peptide they display, we interchangeably use the terms *enriched peptide* and *enriched clone*, while in reality it is the barcode sequencing reads of the antibody bound clones that we measure. When a peptide is enriched, we consider it to be a result of an antibody binding event. An enriched clone is sometimes also called a *hit* or a *binding*. An *IP run* is a set of experimental plates processed together in one laboratory session (Figure 1A). In most cases an IP run consists of 4 or 5 plates. A *Sequencing run* is a set of plates (mostly 7 plates) that is sequenced together in one run of the sequencing machine on one flow cell (Figure 1A).

**Figure 1:**
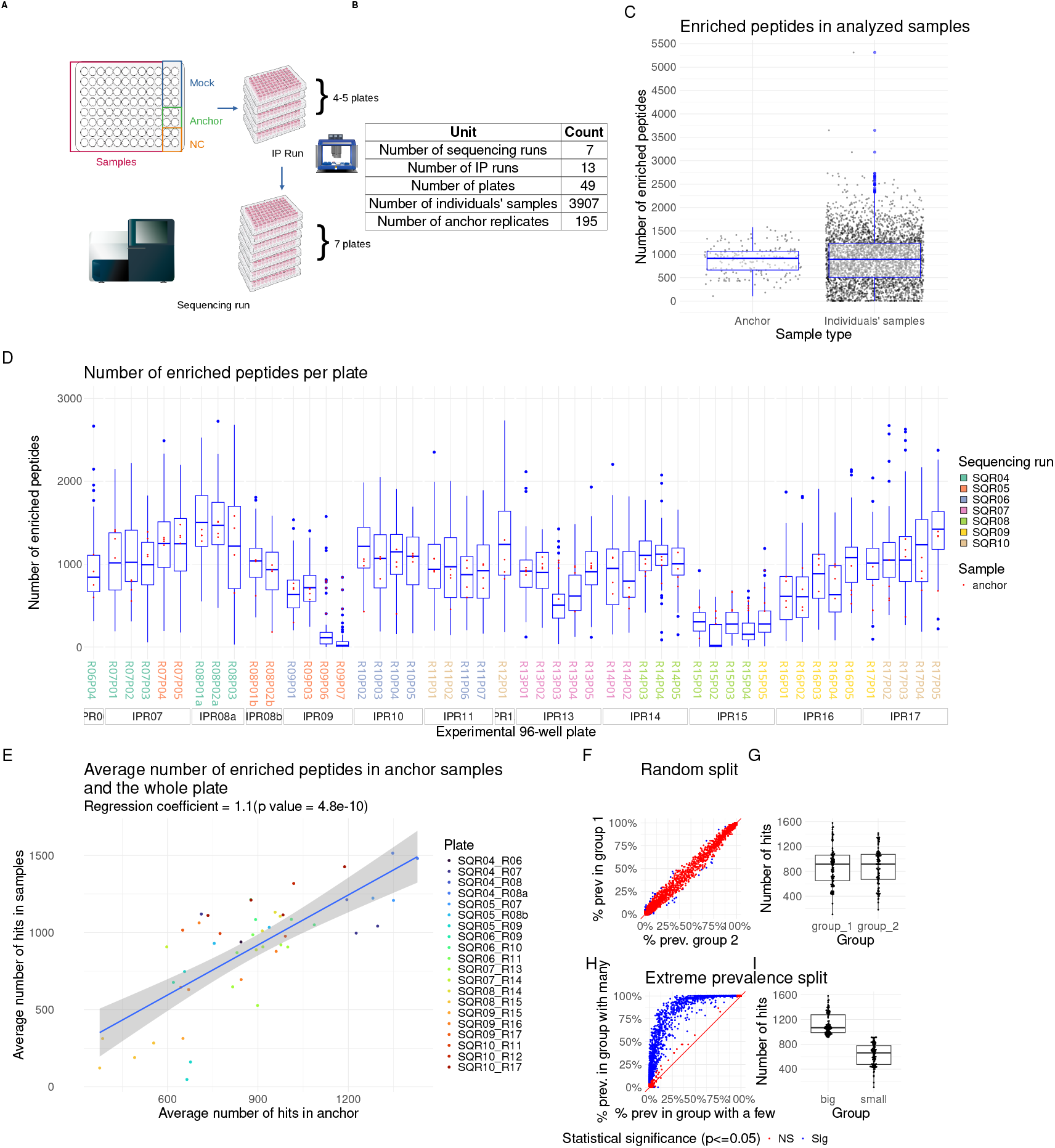
The number of enriched peptides can vary between runs/batches and is to a lesser extent confounded by the experimental plate within the run. **A**: Sample layout on the 96-well plate for all experiments. Each plate contains 80 individuals’ samples, 8 mockIPs (no added antibodies), 4 anchor replicates (a serum pool mixture of 12 individuals) and 4 negative controls. **B**: Overview of the number of samples in this study. Occasionally, the IP reaction fails resulting in the sample count not being a multiple of 80. **C**: The number of enriched peptides per sample varies more than 4-fold across 195 anchor replicates, with individuals’ samples showing even higher spread. The number of hits is therefore not an informative feature for downstream analysis. **D**: The number of enriched peptides on plates from the same IP run tends to be similar while the sequencing run does not seem to be a driver of the batch effects. Plates from the same IP run, but different sequencing run have a similar number of hits to other plates from the same IP run. Each boxplot represents hits from samples on one plate. Anchor replicates are marked as red points. The boxes under the x-axis represent plates from the same IP run, while the color of the plate label represents a sequencing run. **E**: Averaged number of enriched peptides in individuals’ samples and anchor replicates from the same plate show a linear relationship with the slope of almost 1, further confirming similar behavior of samples from the same plate. **F**: A random split of anchor replicates into two groups reveals similar, but not identical prevalence of peptide enrichment per group. Every dot represents an enriched peptide and the position is determined by the relative prevalence of the enrichment in one or the other group. Some of the differences reach nominal statistical significance (p *<*0.05) using chi-square test. More examples of random splits are shown in the Supplementary Figure 5. **G**: Samples from groups in panel F contain a comparable number of hits. **H**: Prevalence of hits in two groups of anchor replicates split based on the number of enriched peptides (more than median and less than median number of hits) shows how an analysis could be confounded by the quality of the PhIP-Seq run. In this extreme case, we observe the characteristic shape revealing that some hits are only detectable if the overall hit number is high. **I:** Groups for panel H were made such that the difference in the number of hits per sample is large while the group sizes are the same.

### Layout

We conduct PhIP-Seq experiments in 96 well plates where 80 wells are for experimental samples and 16 are for controls. Out of the 16 controls, 8 belong to *mockIPs*, 4 to negative controls (*NC*) and 4 to positive controls (*anchor*). The layout is depicted in Figure 1A. MockIPs are immunoprecipitations of the library without serum added and serve as a form of a negative control for background binding of phages or incomplete washing. NC or negative control samples are those where no phages were added and serve to detect contamination. The anchor sample (Figure 2A) is a mixed serum pool of 10 individuals that is added to every plate in every experiment. It serves to detect unusual behavior of a PhIP-Seq run and assess technical variation. The 4 instances of the anchor sample measured on a single plate or anchor sample measured across multiple experiments represent technical replicates.

**Figure 2:**
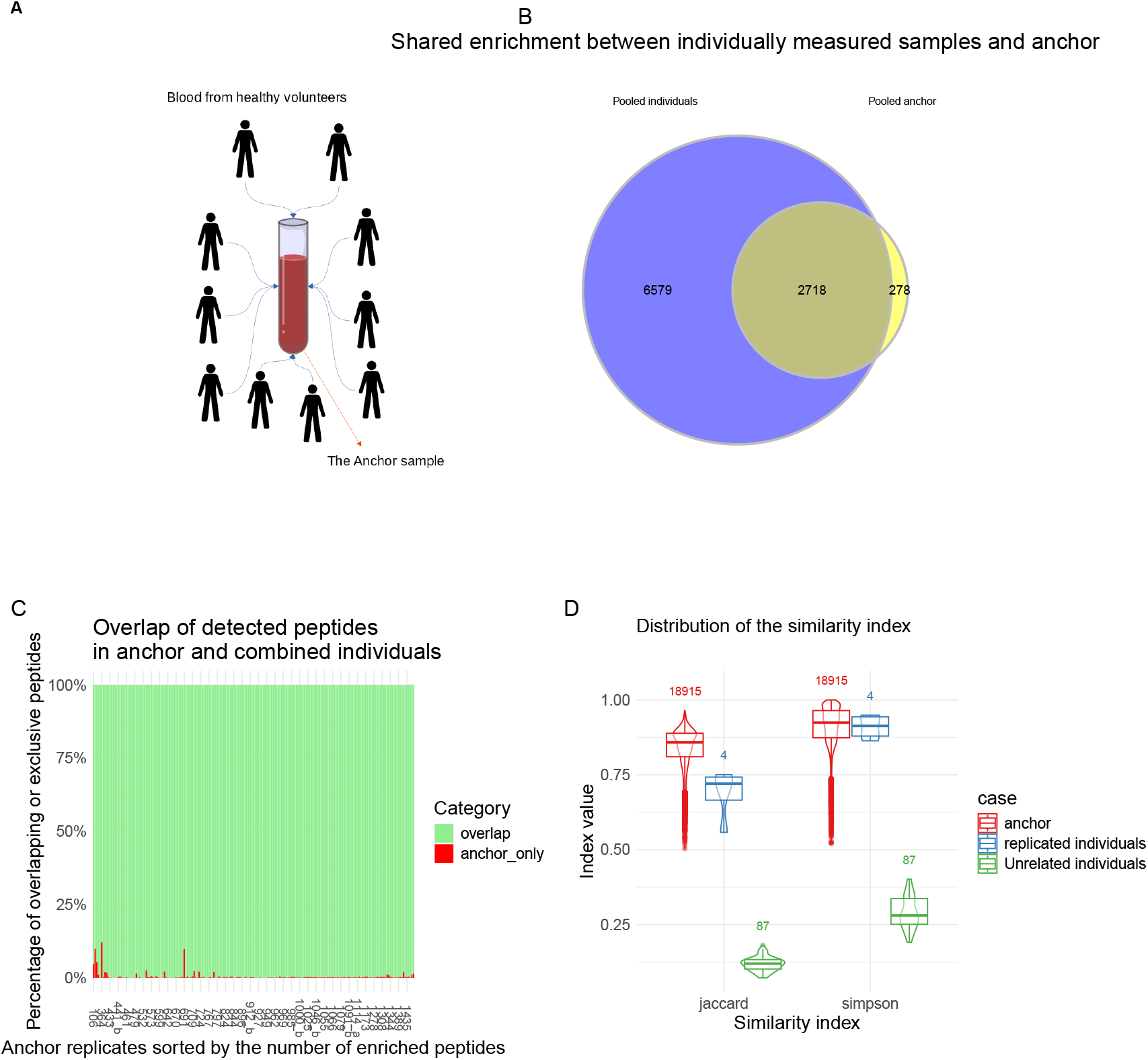
Hits detected in anchor replicates are reliable, with false negatives being the predominant source of error. **A**: The anchor sample is created by mixing sera of 10 healthy donors. Enough is created so that the same serum mixture can be used in many subsequent PhIP-Seq runs. **B**: While the hits detected in anchor are mostly a subset of hits from combined individuals (10 donors), the overall number of different hits from the individual serums is larger than what is detected in all anchor replicates combined, indicating that the serum complexity influences the assay sensitivity. **C**: Hits from anchor replicates mainly overlap the hits detected in individual sera that constitute the anchor, confirming that anchor hits represent true antibody binding events. Each bar shows the relative proportion of hits exclusive to the anchor replicate (red) or hits overlapping with the pooled hits of individual sera constituting the anchor sample (green). Anchor replicates are sorted by the number of hits. **D**: Similarity measured by Jaccard and Simpson indices (Kulczynski index in Supplementary Figure 4) is high between anchor replicates and individual sera measured in duplicate, while for unrelated individual samples it is small, showing that even with the substantial noise in the data, sample-specific information is still captured.

### Enrichment detection

As washing does not completely remove phages not bound by the serum antibodies, it is necessary to use a statistical model to detect bound phages via enrichment of their reads in the sequencing set. Various methods have been applied for enrichment detection [4, 25–27]. We use enrichment detection by fitting a generalized Poisson distribution on the counts of the phages that had the same initial count in the library (same input count) and call phages enriched if the p value of the count is less than 0.05 after Holm correction for multiple testing [28]. The normalization procedure follows the description in ref. [4].

### Analysis of cohort PhIP-seq data

The analysis commonly consists of the comparison of the antibody repertoires of healthy individuals and patients. The groups should be age and sex matched as those factors influence the antibody repertoires [11]. Previous analyses are described in [15, 16, 18, 29].

### Analysis of the anchor sample

The anchor sample is processed 4 times per plate and is included in every run. We have collected 195 technical replicates of the anchor sample (sequencing of one sample failed) which allows us to study the replicability of the PhIP-Seq assay (Figure 1B). Additionally, we ran PhIP-Seq with 10 individual’s sera whose mixture constitutes the anchor sample.

### Control replicates of patient samples

In addition to the anchor sample and individual sera that constitute it, we have measured 76 serum samples in our lab (Medical University of Vienna, Vienna, Austria) and compared them to the results published in the past [30, 31] (Weizmann Institute of Science, Rehovot, Israel). The phage library was identical by design in both experiments (produced from the same glycerol stocks), but the phages were created separately by expanding them to a large quantity from the glycerol stocks by plate amplification (as described by Mohan et al. [24]), and it differed in composition due to random factors in the phage production and expansion step. Computational processing for both datasets was identical. For evaluation of the effect of the biological material used for PhIP-Seq reactions to the results, we conducted PhIP-Seq experiments on another set of 113 individuals, but measuring both plasma and serum from the same individuals.

## Results

In our PhIP-Seq implementation, experiments are conducted on 96-well plates, each containing 80 individuals’ samples and 16 controls: 8 mockIPs (immunoprecipitation without added serum), 4 negative controls (neither phages nor serum added) and 4 anchor samples. The anchor sample is a serum pool from 10 healthy individuals included in quadruplicate on every plate. Over the course of this study we collected 195 such anchor replicates from 49 experimental plates measured in 13 runs, allowing us to characterize technical variability and replicability independently of any individuals’ samples to be measured. A peptide is considered enriched, or a hit, if its read count after immunoprecipitation is higher than expected under a null model (generalized Poisson distribution, p*<*0.05 after Holm correction), out of a total library of 357 192 peptides. In our discussion we use the term enriched peptides instead of bound peptides to emphasize that we are discussing the properties of the PhIP-Seq method rather than the biological meaning.

### Enriched peptides in replicates

#### Number of hits

Important information about the PhIP-Seq data is the replicability of the results, i.e., how much will the results differ between runs. Differences in the set of enriched peptides between the replicates stems from a different number of false positive (FP) and false negative (FN) hits.

The number of enriched peptides varies substantially for the same anchor sample measured multiple times (Figure 1C,D,E). 195 technical replicates of the anchor sample vary in the number of enriched peptides more than 4-fold, while the variability among individuals’ samples is even larger, but the median number of hits is comparable to the anchor sample (around 950) (Figure 1C). We also observed that individuals’ samples and anchor sample replicates processed on the same 96-well plate behave similarly in terms of the number of hits (Figure 1D). Regression of the average number of hits among individuals’ samples from a plate and the average number of hits from associated anchor replicates has a coefficient of 1.1 (Figure 1E). On an aggregated level of the IP run, plates from the same run usually all show either increased or decreased number of detected hits, while the sequencing run does not seem to be a large driver of batch effects (Figure 1D, R15P05 is sequenced in the sequencing run 9, but behaves similarly as other plates from the IP run 15 that are sequenced in the sequencing run 8).

#### Identity of hits

Not all peptides have the same probability of being enriched. When anchor replicates are randomly divided into 2 groups (Figure 1F,G) the prevalence of hits in the two groups is similar and a few peptides are nominally significantly differentially prevalent (chisquare test). However, different random groupings could give different number of significantly different peptides (Supplementary Figure 5). On the other hand, when anchor replicates are split into groups that maximize the difference in the number of hits (top 50% of replicates with the largest number of enriched peptides vs. bottom 50%, representing the worst case, Figure 1G,H), it becomes clear that some peptides are more readily detected when the number of hits is in general higher as their prevalence in the group of samples with more hits is larger. The apparent difference in the antibody repertoire between the two groups could be reduced by increasing the number of samples per groups to compensate for the random factor in the retrieved data as the variability in the binding prevalence drops quickly once the sample size per group exceeds 250 for the comparison where we do not expect differences (Supplementary Figure 3).

**Figure 3:**
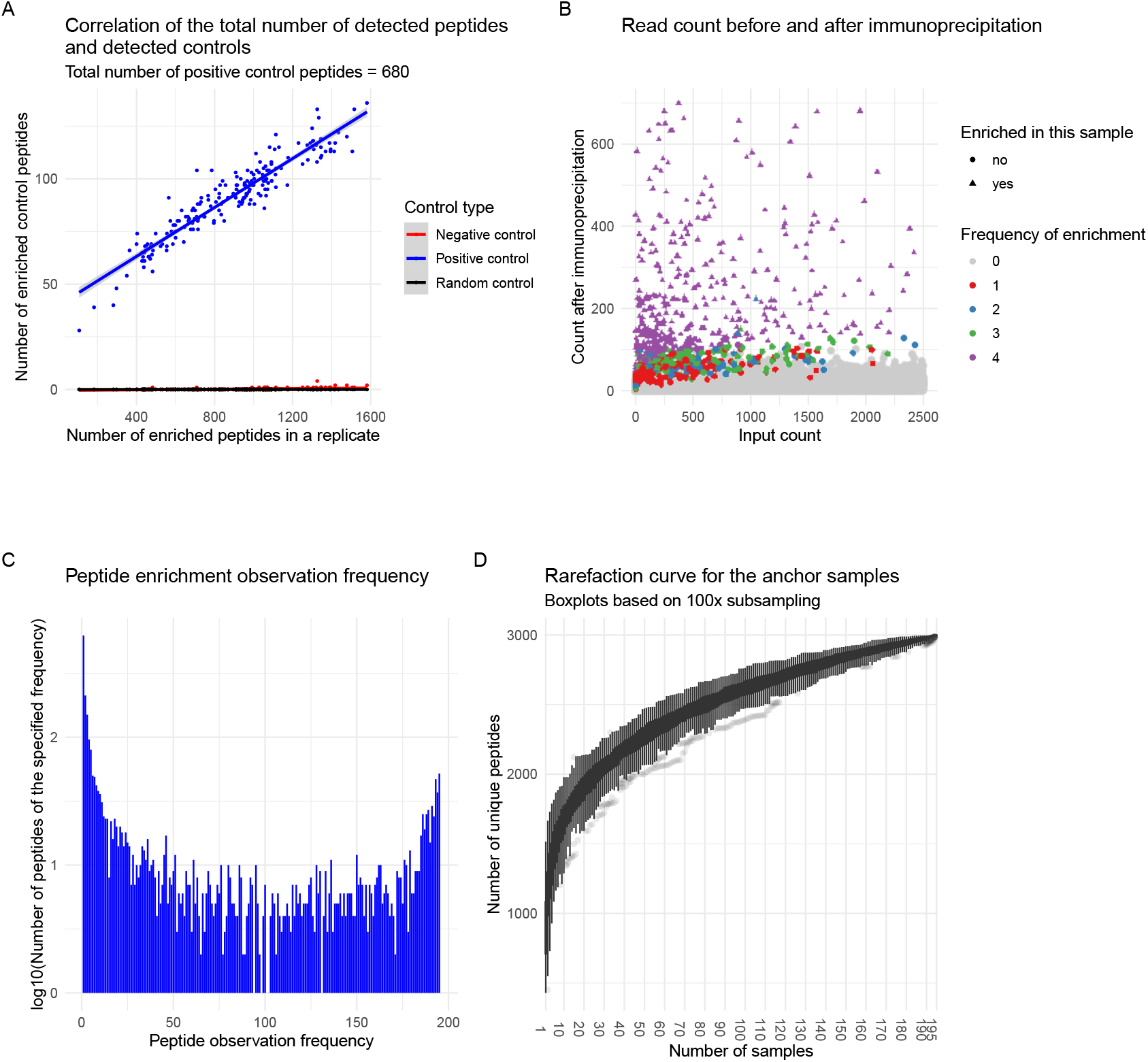
Technical replicates increase the comprehensiveness of detected antibody repertoire. **A**: The number of enriched positive control peptides increases with the total number of enriched peptides per anchor replicate, while the number of enriched negative control peptides remains near zero. Additional peptides detected in high-count replicates therefore represent true detections rather than increased false positives. The figure shows the data for all 195 anchor replicates. **B**: Input read counts versus immune-precipitation read counts for one anchor replicate, with points colored by how frequently each peptide was enriched across all four replicates on the same plate. Some peptides enriched in other replicates have counts indistinguishable from background (non-enriched phages) in this replicate. However, strongly enriched clones tend to be enriched in all replicates. This demonstrates that failure to detect peptide enrichment arises from the stochastic variation in immunoprecipitation. **C**: Combined peptides from all anchor replicates are observed at different frequencies. Two peaks around 100% and 0% point to easy-to-detect peptides and peptides that a single experiment cannot reliably detect. Notably, the shape of the histogram is similar to observed peptide frequencies in the human population [11]. **D**: Rarefaction curve based on 195 replicates does not saturate which is consistent with the Venn diagram in Figure 2 C. For the anchor sample, more replication is needed to reach saturation. As the anchor sample is a mixture of 10 sera, curve from individual sera would likely saturate at a different rate.

#### Quality control of hits

As established earlier, technical replicates produce different number of hits per sample. It is important to verify if the difference in hits stems from retrieval of the real antibody binding events. Our PhIP-Seq library contains various peptides labelled as positive (antibody binding reported in the literature, even with PhIP-Seq [11]) or negative controls, and in case a larger number of hits corresponds to the enrichment due to antibody binding, we should observe an increase in the number of positive control peptides among the hits without an increase in the number of negative control peptides. Figure 3A shows a trend where the number of retrieved positive control peptides increases linearly with the number of enriched peptides in anchor replicates, while the number of negative controls (together with random peptides) does not increase. To show that the difference in the number of hits originates from the efficiency of the immunoprecipitation and magnetic-bead pull-down, Figure 3B and Supplementary Figure 2 show the read counts before and after immunoprecipitation for one anchor replicate and highlights the peptides enriched in other replicates from the same plate. The read counts of the peptides missed in the depicted replicate are not different from the background phages. Importantly, peptides have a different probability of being enriched assuming the corresponding antibody exists in the sample which is depicted in Figure 3C. A similar distribution is also observed in the prevalence of the hits in the healthy population [11]. Interestingly, two peaks at the borders show that still some peptides are almost always detected (right peak) and others are singletons even among 195 replicates (left peak). To retrieve all possible hits, enough technical replicates are required, depending on the complexity of the sample. However, for a highly complex sample such as the anchor sample (mixture of 10 individuals), not even 195 replicates are enough to reach the saturation of the rarefaction curve (Figure 3D).

### Validity

It is not realistically possible to obtain a gold standard for antibody binding data at scale with PhIP-Seq. Besides practical limitations of conducting hundreds of thousands of ELISA assays, a conceptual problem with a ground truth is that with PhIP-Seq, it is theoretically only possible to detect antibody bindings against linear epitopes and conformational epitopes in case the displayed peptides fold correctly. To evaluate the validity of the antibody binding data from PhIP-Seq, we compare the hits from the anchor replicates and the individual sera that constitute the anchor sample (Figure 2A). The practical reason for this is that our anchor sample is a pool of several individuals. Since we want to keep using the same anchor sample also for future work, we need a large quantity. Rather than collecting multiple blood draws from a single person, we hence collected blood from several individuals and mixed them into a pool. Overlap in the hit sets between the pooled individually measured sera and the anchor replicates is expected. Hence, we can compare individual measurements of the 10 samples that make up the anchor pool to the results of them together combined in the same pool. Thereby we can assess the recovery of the assay, if single individuals’ reactivities all appear in the anchor. Figure 2B shows that the hits from the anchors are a subset of hits obtained from the individuals. Together with the fact that we obtain similar number of enriched peptides in anchors and patient samples (Figure 1C, Supplementary Figure 6), the data suggests that there is a plateau of the number of hits we can detect with the current setup: While we detect about 1000 peptides per individual sample of the 10 people that make up the anchor pool, we do not detect 10 000 peptides in the pooled anchor sample, but rather also around 1000 peptides (this calculation is simplified and ignores that multiple individuals would have antibodies against the same antigens such as vaccinations and common infections, although, overall about 9200 peptides are detected across the 10 individuals measured separately, see Figure 2B). This finding suggests, that there is a technical limit in our PhIP-Seq implementation, potentially owning to aspects such as the amount of magnetic beads used for the IP pull-down.

Besides the aggregated hits of anchor replicates, we also evaluated the overlap of the hit set of the individual replicates against the pooled hits of individual sera (Figure 2C). In general, anchor replicates contain only a few hits that are not also present in individually measured sera and the number of those hits does not increase with the number of enriched peptides per replicate.

### Replicability

#### Similarity of replicates

With any method, knowing how reproducible the results are (for reproducibility and replicability, see ref. [32]) directs the downstream analyses and limits possible conclusions. To estimate the similarity, Jaccard and Simpson similarity indices were calculated for all pairs of the replicates of the anchor sample and visualized the distribution of both (Figure 2D). In the ideal case, the pairs of replicates would have both indices equal to 1. The actual values are lower, with Jaccard index being even smaller. Given the variation of the number of hits, it is expected the Jaccard similarity is smaller, but the value of the Simpson’s similarity index *<*1 means that the set of obtained hits is a partially random subset of all possible hits. This is in accordance with the observation about the hits present in all and in a few of the replicates (Figure 1B, Supplementary Figure 1). Pairs of unrelated samples (individuals contributing serum to the anchor sample) have much lower similarity index values, which is expected since the antibody repertoires are largely individual [11]. The result shows the expected similarity between the replicated samples and points out that the selection of the similarity metric is crucial for comparing samples. However, even when using the Simpson’s similarity index which takes into account the unequal number of peptides, the value is not 1 for technical replicates.

### Effect of a biological material - serum vs. plasma

Blood plasma, the antibody-containing fraction of blood, can also be sampled as serum by removing coagulation factors. Ideally, measuring serum or plasma would not matter for a PhIP-Seq experiment as the antibody content of both should be the same. However, we have observed peptides that are preferentially enriched either in serum or in plasma samples (Figure 4A). In serum samples, we observe enrichment of peptides originating from fibronectin-binding proteins while in plasma samples those are mostly depleted. On the other hand, we observe enrichment in plasma samples and depletion in serum samples of peptides originating from staphcoagulase and fibrinogen-binding protein (Figure 4B). We propose that this effect could be explained by the interaction of the peptides retaining the function of their original protein (for example fibronectin-binding protein) and the coagulation factors (such as fibrinogen) which blocks the antibody binding to the displayed peptide (Figure 4C, D). Those proteins are part of the library because they appear in the Virulence Factor Database (VFDB) [33]. Via interaction with the coagulation factors, bacteria use them for attaching to the host tissue and evade immune response [34].

**Figure 4:**
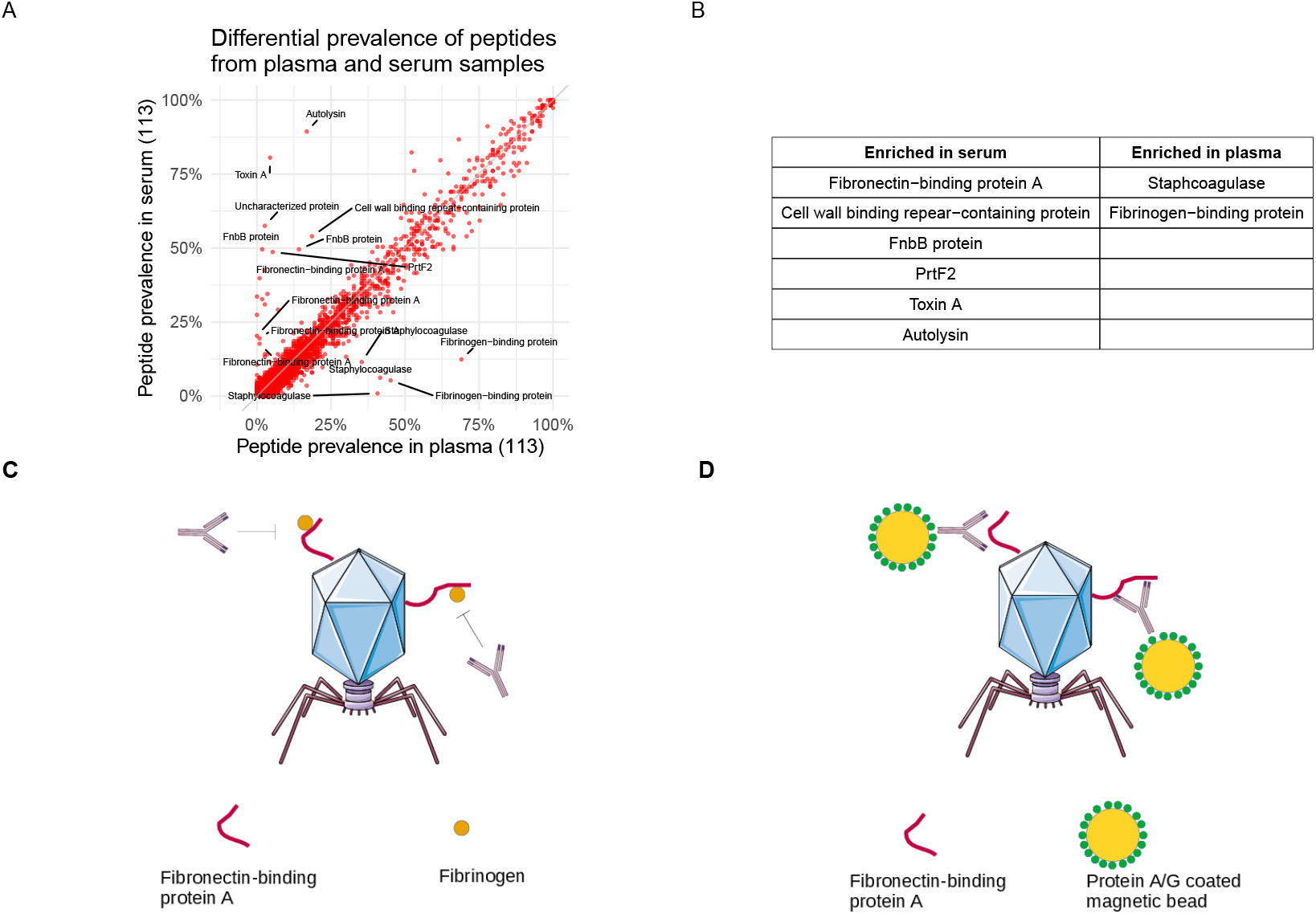
The type of biological material (serum or plasma) influences PhIP-Seq enrichment profiles, likely through interactions between phage-displayed peptides and plasma proteins. We evaluated the effect by measuring both plasma and serum samples for 113 individuals. **A**: While most of the peptides show similar prevalence regardless of being detected in plasma or serum samples, a number of peptides mostly originating from Fibronectin-binding protein A from *Staphylococcus aureus* are more readily detected in serum. In contrast, peptides from Staphcoagulase are more often detected from plasma samples. **B**: Some antibody bindings to peptides derived from certain proteins are more readily detected in serum and some in plasma of the same individuals. **C**: In plasma, coagulation factors are available for binding to peptides from Fibronectin-binding protein A which block potentially present antibodies from binding the peptide. D: Coagulation factors are absent from serum and antibodies are free to bind phages expressing peptides from Fibronectin-binding protein A and can be subsequently enriched by pull-down with magnetic beads.

**Figure 5:**
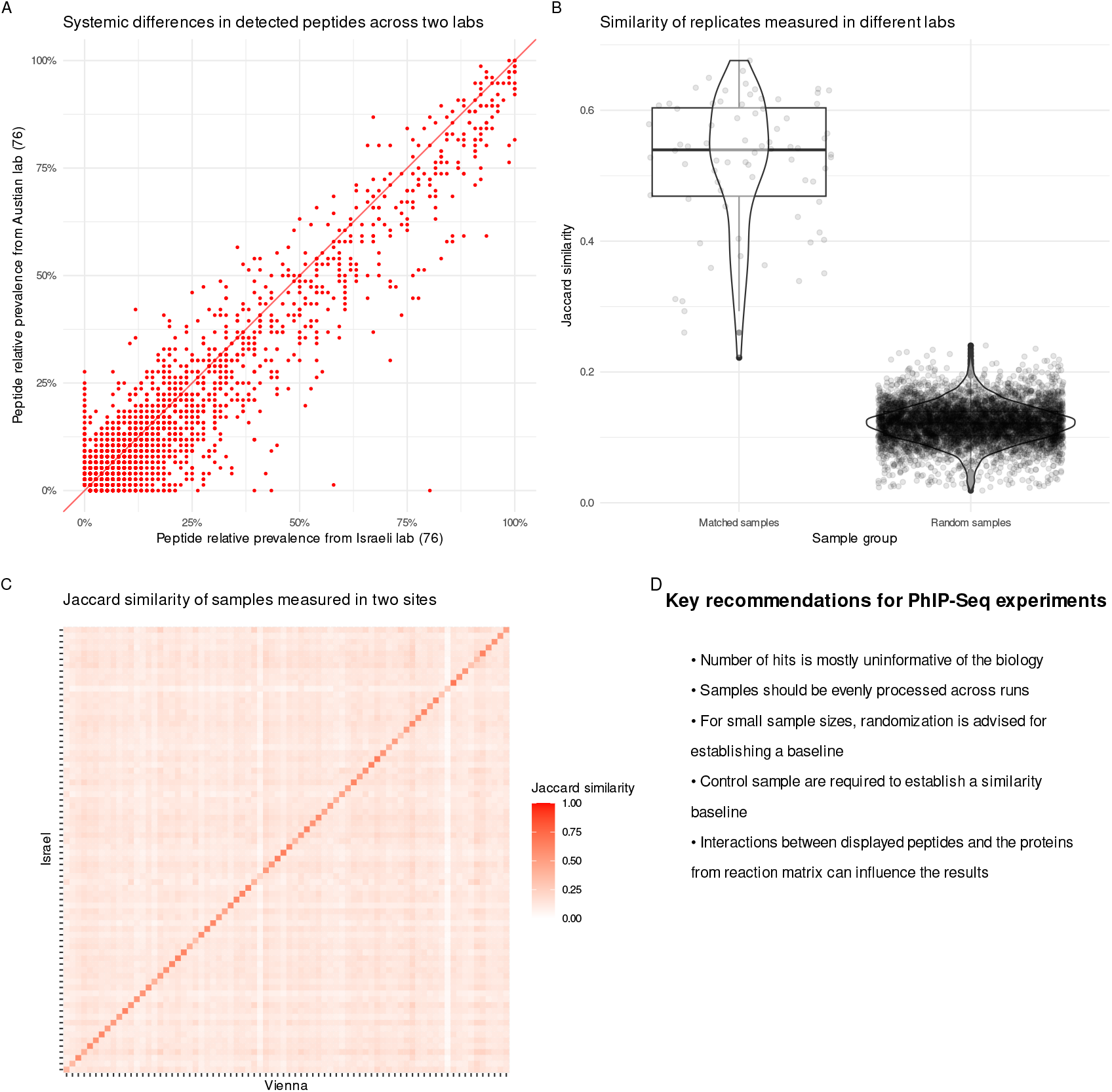
PhIP-Seq experiments in different laboratories yield comparable enrichment profiles with slight site-specific differences. **A**: Most peptides have similar population prevalence regardless of the lab where PhIP-Seq was conducted, but there are peptides that are never detected in one or the other lab (dots lying on x and y axes). As libraries were the same in design and the experimental protocol was similar, it seems that the library expansion (which was done separately in different labs) influences the ability to detect certain peptides. **B**: Even with differences in the library composition originating from randomness in phage expansion, matched samples still have larger similarity index than what is observed between unrelated samples. **C**: Jaccard similarity between matched and all other pairs of samples in the heatmap form. Matching of the PhIP-Seq profiles between the corresponding samples is depicted by a diagonal of the heatmap and standing out. **D**: Summary of the recommendations for approaching PhIP-Seq experiments and data interpretation.

### Cross-laboratory replication

PhIP-Seq is an emerging technology and the laboratory where experiments are conducted could influence the outcome. We analysed the results of samples measured in two laboratories using the same glycerol stocks of clones of phages and looked for systematic differences in the retrieved enrichment profiles. The two experiments are different also in the machine used for sequencing (Illumina NextSeq in Israel vs. Illumina NovaSeq 6000 in Vienna) and the liquid handling robot (Tecan Freedom Evo in Israel vs. Tecan Fluent 1080 in Vienna). Most of the peptides show comparable prevalence among samples in both laboratories, but there is a number of peptides that are exclusively detected in only one or the other lab (Figure 5A). Even though the libraries have a slightly different composition due to stochastic factors in library expansion, the difference is not enough to explain the difference in the results obtained in both labs (Supplementary Figure 7). However, even with systematic differences in the enriched peptides, the similarity analysis shows that we are able to match samples originating from the same individual almost perfectly (Figure 5B, C).

## Discussion

### Antibody binding recovery

We have shown that the peptide bindings obtained by PhIP-Seq as implemented in our laboratory are trustworthy, but at the same time not always recovered to the same extent in repeated experiments, which has important implications for downstream data analysis. It was already stated by the inventors of PhIP-Seq that absence of the signal does not mean the absence of the corresponding antibody [4], but the extent of the issue was not quantified, to the best of our knowledge. It is worth noting that the authors of the original study and other studies about autoimmunity may expect only a limited number of clones to be enriched, while we expect with our microbial antigen library a couple of thousands of enriched clones per sample. In a study on the antibody response against diverse viruses, Xu et al. have shown by ELISA that false negatives are a major drawback of the method [9]. Our work partially quantifies the extent to which clones that could be bound by the serum antibodies are detected or missed. Importantly, there seems to be an upper limit of the number of enriched clones which we can detect in a single run. To detect more, technical replicates are needed, however, even when re-measuring a single serum sample 195 times, we did not reach saturation in the number of detected clones.

The anchor sample measured as a technical replicate consisted of a mixture of 10 individual sera and is not completely representative of the samples where a single serum is measured. Due to mixing, the concentration of the antibodies that target the same peptide is amplified and the concentration of the low abundance antibodies is reduced, which makes it more difficult for them to bind the phage displayed peptides and cause enrichment. To detect full binding capacity of a serum sample, it is necessary to measure it multiple times. For analyses involving a small number of samples, measuring the serum samples in replicates might be advisable. For comprehensive characterization of a single sample, we suggest making a rarefaction curve to estimate the completeness of the recovered bindings. From our experiments, we are not able to derive a simple estimate of the false negative rate, as it would not be transferable to other experimental settings. In addition to the anchor being a mixture, the flexibility in the enrichment calling threshold, and the lack of a gold standard, the false negative rate also depends on the library composition and is therefore not universal. It is worth noting that some protocols make use of multiple rounds of phage amplification after IP [35] or suggest pre-washing of the serum with wild type phages [36]. Such procedures may influence the recovery of the bound phages.

### Alternative detection methods

It must be noted that there are alternative statistical methods to detect enriched clones besides modeling of the output counts as a generalized Poisson distribution (GP). A small improvement over the GP method is to fit the zero-inflated version of the generalized Poisson distribution (ZIGP) to the data [9]. Adapted from the field of RNA-seq are EdgeR [25] and BEER [26], the two methods applied to PhIP-Seq data that compare the sequencing counts of the clones of the samples to counts obtained from mockIPs. The Z-score method also compares the sample counts to the counts from mockIPs, but aims to be easily understandable and interpretable [27]. Our data collection is obtained with libraries described by Vogl et al. [11], Klompus et al. [22] and Leviatan et al. [12] where the GP method is used to obtain enrichment. To ensure comparability, we have continued using the GP method to obtain the enriched peptides.

### Statistical model adjustments

It is however possible to modify the GP method in various ways, most obvious being changing the p-value threshold for calling enriched peptides. The decision ultimately depends on the acceptable false positive (FP) and false negative (FN) rates. However, as it is practically impossible to precisely measure the antibody reactivity against all 357 192 peptides to obtain the gold standard for a serum sample and calculate the true FP and FN rates, we kept the threshold at p *<*0.05 after Holm correction for multiple testing [28].

### Factors influencing the experiment

For each particular PhIP-Seq implementation, there are specific factors that influence the experiment results. Enrichment detection methods assume the majority of the clones is not enriched to easily model background number of reads. For better enrichment detection, a library should be large and uniform (equal abundance of the clones in the stock) and only a minority of the clones should be bound by any serum. Those factors are in control of the experimenter setting up a PhIP-Seq experiment. Factors outside the experimenter’s control are serum antibody composition, antibody-antigen binding strength and antibody specificity or cross-reactivity. Low antibody concentration, weak binding strength and cross-reactivity reduce the enrichment of the clone displaying a target peptide and increase the chance of not detecting it.

### Effect of the biological material

An interesting observation of a systematic difference between the enrichment profiles obtained from serum or plasma samples partially works in favor of validating PhIP-Seq as a sensitive method. Both serum and plasma are expected to contain identical antibodies, but the binding of the antibodies and their targets can be influenced by the presence of any other molecules in the reaction well. The difference between serum and plasma is in the content of coagulation factors including fibrinogen and prothrombin, which serum lacks. Fibronectin-binding protein A binds fibronectin and fibrinogen [37], which are present in plasma and not in serum. Due to binding of the peptide expressed on the phage to proteins, the interaction with the antibody is prevented, and the clone is not enriched. On the other hand, serum lacks fibrinogen and the expressed peptide originating from fibronectin-binding protein A is free to interact with the corresponding antibody causing the subsequent enrichment. However, fibrinogen-binding protein is also supposed to bind fibrinogen in plasma and hinder the interaction with the corresponding antibody, but we observed enrichment of clones carrying peptides from fibrinogen-binding proteins only in plasma samples. Similarly, staphcoagulase binds prothrombin [38] which is also present in plasma and not in serum, making it unusual that we observe enrichment preferentially in plasma samples. We are not able to entirely explain the effect, but these findings suggest that the peptides expressed on the phage surface can still keep some binding properties of the protein they originate from and which consequently influences the results of the immunoprecipitation reaction by bindings to various serum/plasma proteins and possibly to each other. We have previously demonstrated such effects for protein A/G and gut microbiota antibody binding proteins (Figure 2 from ref. [11]). As PhIP-Seq could be conducted on antibody-containing biological material such as cerebrospinal fluid [6, 39] or aqueous humor [40], a researcher should be mindful of the potential interactions of the matrix containing proteins and the phages from the library. The potential for interaction of the displayed peptides and the matrix/antibodies depends on the library. We envision that the interaction potential for the library of the human proteome would be larger due to the fact that human proteins more often interact with other human proteins than microbial proteins.

### Considerations for the downstream analysis

Given the data we have presented in this paper, we can offer some recommendations for design of the PhIP-Seq experiments and the downstream analysis of the data from PhIP-Seq experiments (summary in Figure 5D):

1. The antibody bindings deduced from enrichment do not necessarily represent a comprehendsive set of possible bindings from a serum. Replicates are recommended when comparing groups with small sample sizes. Consequently, the number of hits should be shown to rule out batch effects between groups of interest, and caution should be applied when using it as a feature in data analysis.
2. The number of enriched clones can show a batch effect dependent on the IP run and to a lesser degree depending on the experimental plate within a run. To minimize the batch/plate effects on the data analysis, plating the samples of the same group on the same plate and measuring them in different runs/batches should be avoided (i.e. putting all cases of a disease on one plate, and all healthy controls on a different plate, and running these two plates in different batches is problematic). Measuring plates with samples from distinct groups in the same batch appears feasible, but potential differences between plates need to be assessed with technical replicates/internal standards (what we call anchor samples). To also rule out this effect, samples from different groups can be distributed on the 96-well plates randomly.
3. For experiments with small sample sizes, spurious differences between the groups could arise due to technical variability. Use randomization to establish a baseline of the peptide prevalence differences between groups.
4. Use enough control replicates (anchor) to estimate a baseline for the sample similarity and detect issues related to a 96-well plate.
5. While interpreting the data, interactions of the expressed peptides with other proteins present in the sample and with each other have to be considered. The opportunities for interaction depend on the library composition.

## Acknowledgements

We would like to thank Melanie Prinzensteiner, Nikolas Basler, and Michaela Fehringer for fruitful discussions. Computational results of this work have been achieved using the Life Science Compute Cluster (LiSC) of the University of Vienna. T.V. gratefully acknowledges support from the European Union (European Research Council (ERC) Start Grant EarlyMicroAbs (project number 101075733) and Horizon Health consortium “ID-DarkMatter-NCD” (PN: 101136582)), as well as support from the EMBO YIP program (#6165).

