## Supplementary figures for "Pitfalls in understanding PhIP-Seq data: technical variability, replicability, and best practices for interpretation"

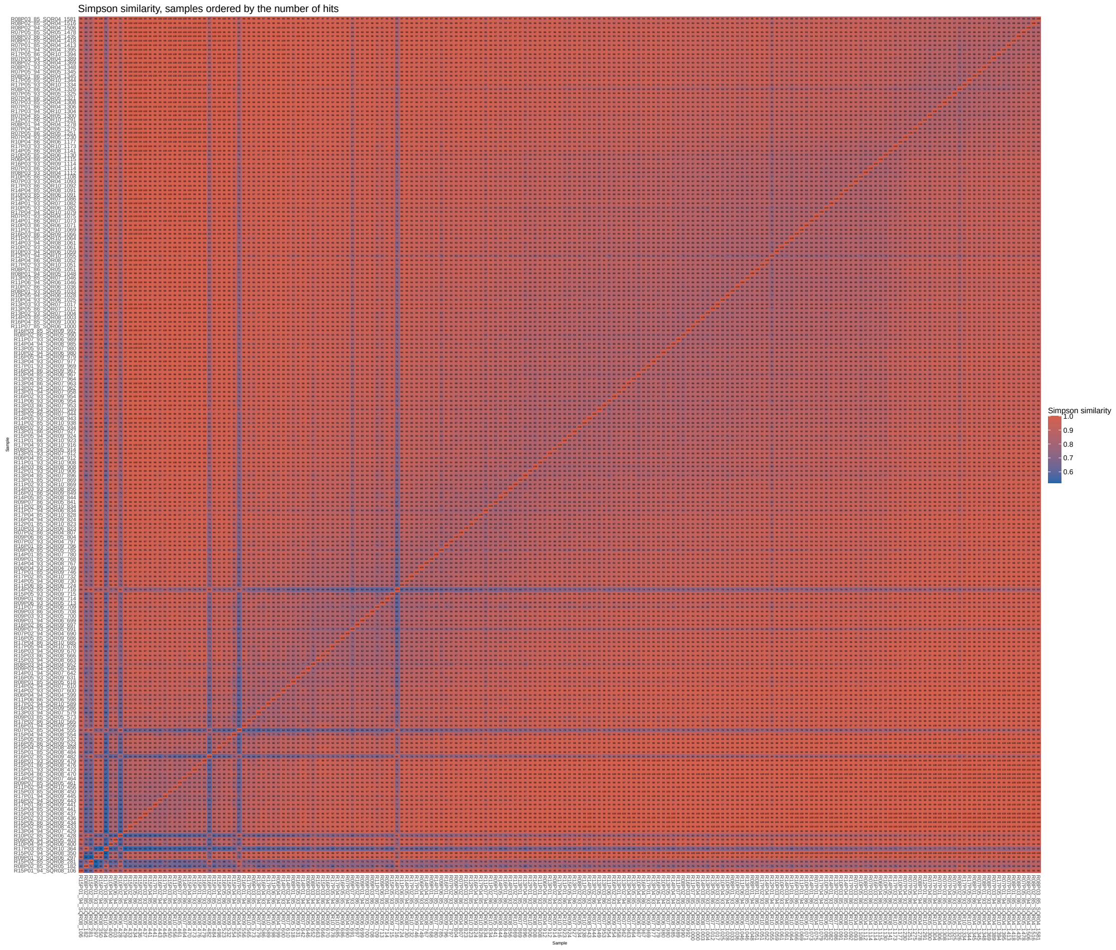


Supplementary Figure 1: Simpson similarity index for all replicates of the anchor sample, sorted by the number of enriched peptides. The last number in the sample label represents the number of enriched peptides in the sample. The heatmap shows the general trend where the replicates with less enriched peptides are less similar to each other than to the replicates with more enriched peptides. The trend suggests that the enriched peptides per replicate are partially a random subset of the all possible enrichments given the serum antibody composition.

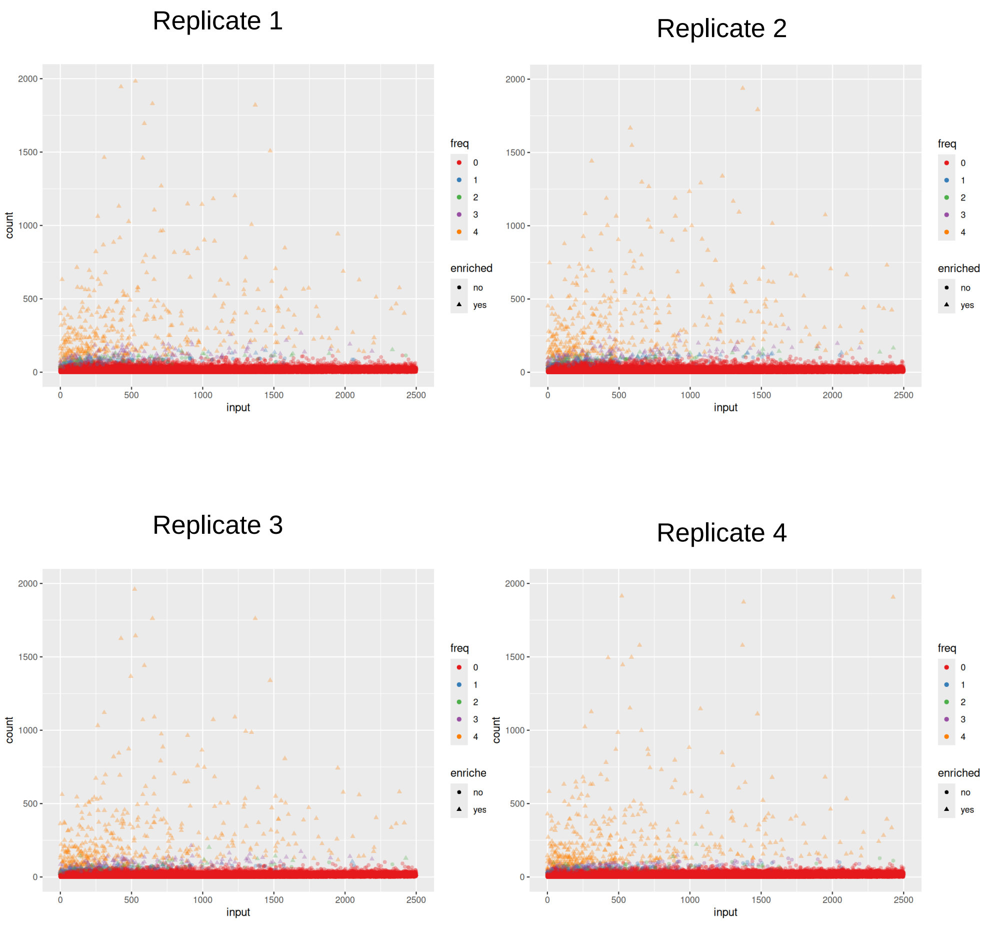


Supplementary Figure 2: Scatter plots showing the input count vs. count after IP. Each point represents a clone and each subplot represents a replicate of an anchor sample from the same plate. The replicates contain the same serum, but the sets of enriched peptides do not overlap 100% between the replicates. The point color represents in how many out of 4 replicates was a certain clone detected as enriched. We can observe that some peptides with enrichment frequency $<4$ are buried within the noise in replicates where they are not enriched. In those cases, it is impossible to detect the binding via enrichment unless the number of false positives drastically increases.

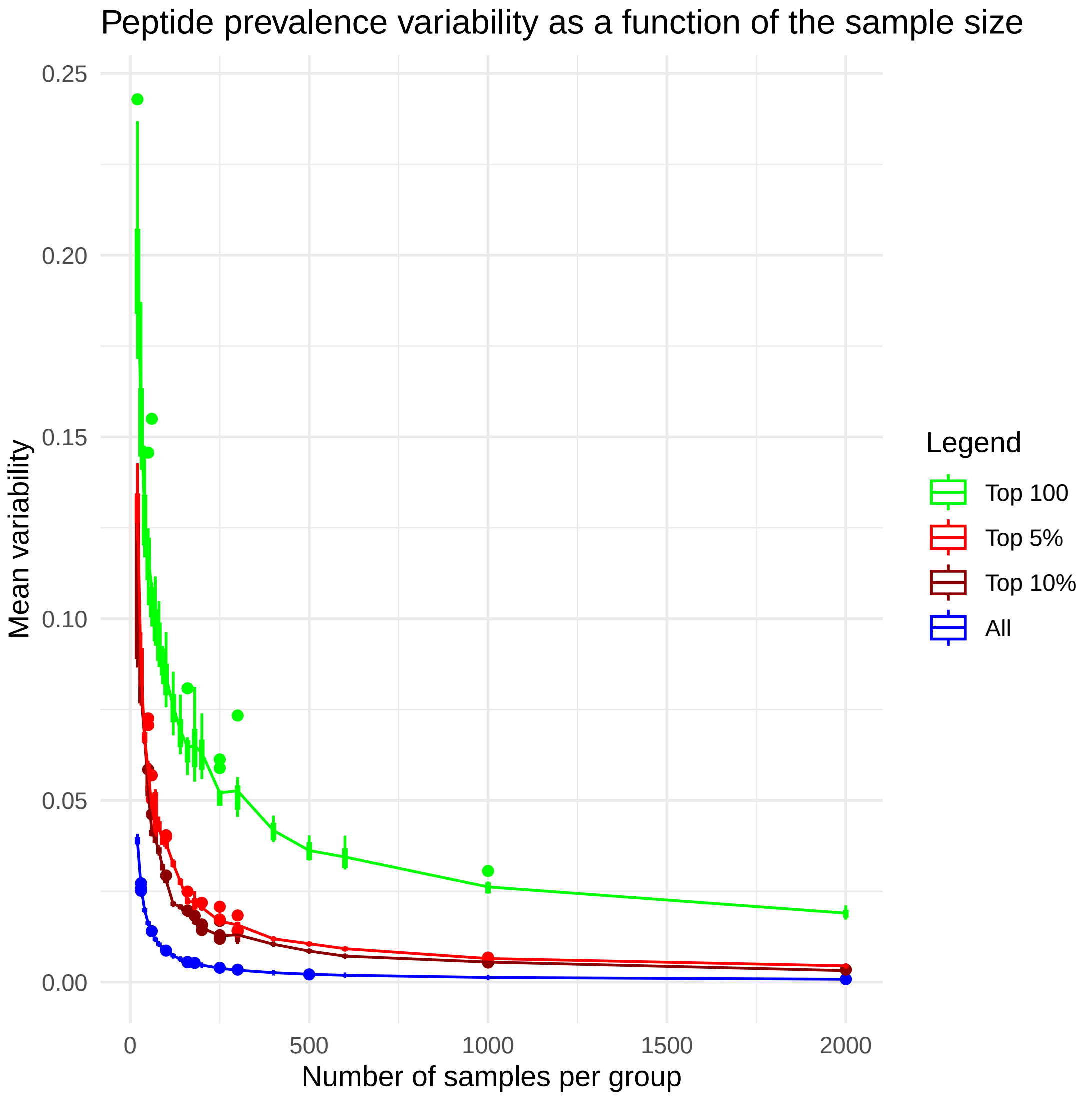


Supplementary Figure 3: Dependence of the hit prevalence variability on the number of samples per group. The groups are made by randomly selecting a number of samples from our collection (size ∼4000). The mean variability is defined as the mean euclidean distance of the peptide to the center line (equal prevalence for both groups). As the most variable peptides are usually the most interesting, the plot shows the variability for different number of most prevalent peptides (all, top 10%, top 5% and top 100). The variability is dropping steeply for sample sizes smaller than 500. Ideally, the group sizes would not be smaller than 500 samples, but the cost of obtaining cohorts this large is usually prohibitive.

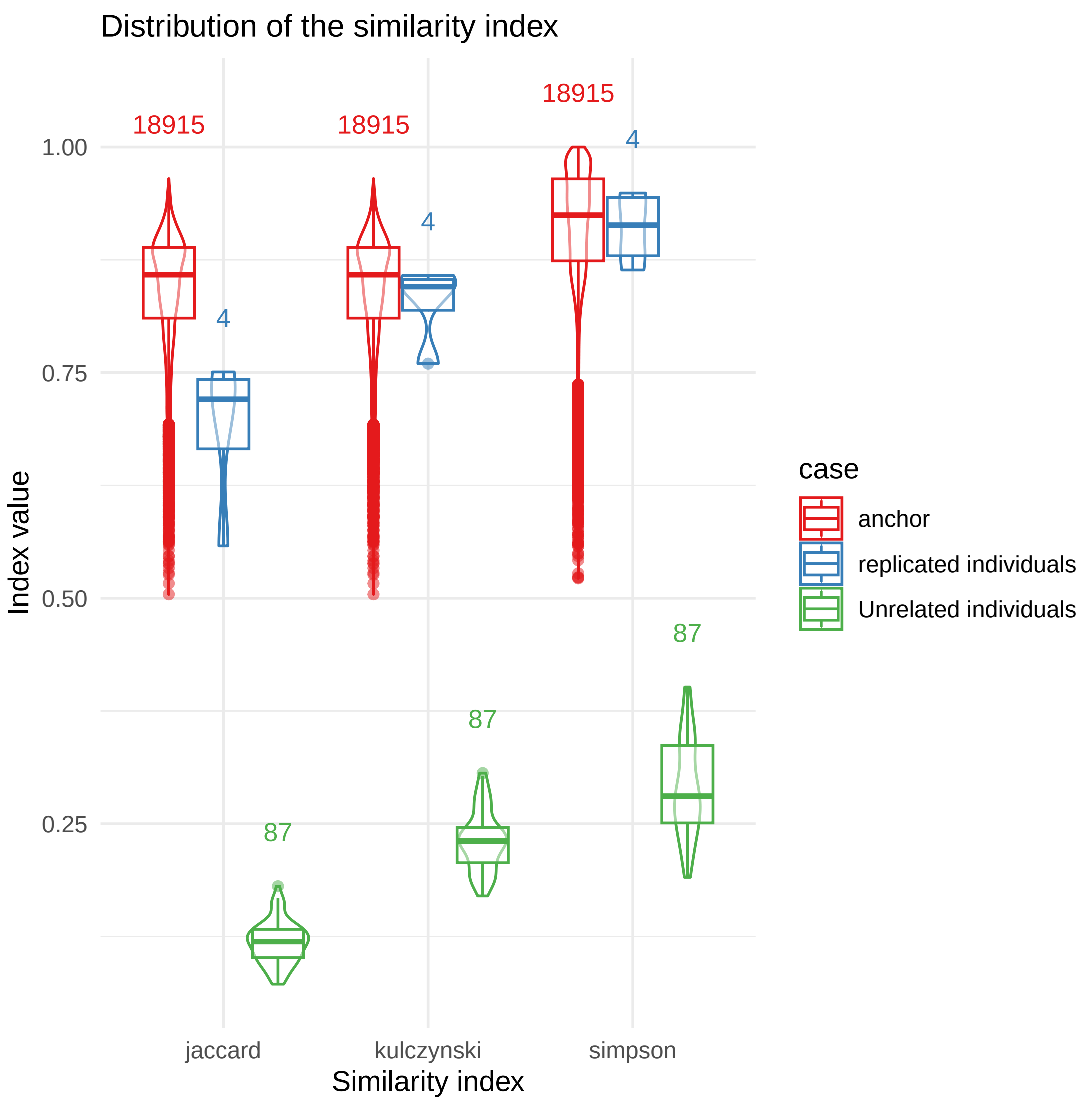


Supplementary Figure 4: Similarity between anchor replicates, replicated individuals and unrelated individuals. As sometimes Kulczynski similarity is used to compare the composition at two sites, we included it alongside Jaccard and Simpson similarities for comparison.

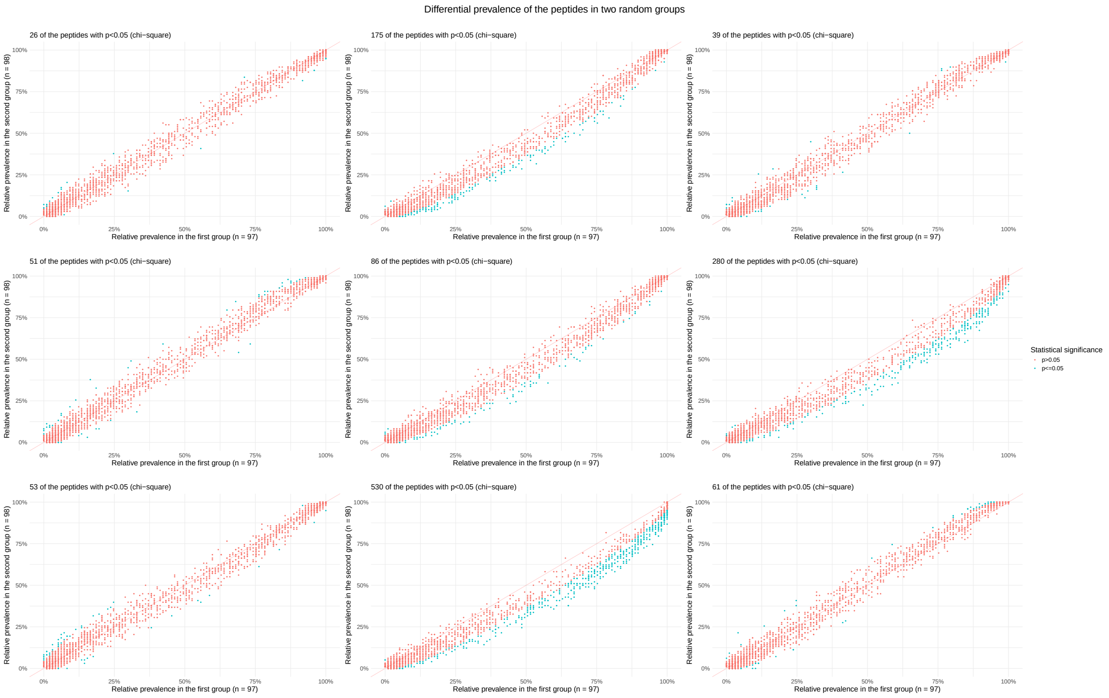


Supplementary Figure 5: Differential prevalence of the peptides in groups made by randomly splitting technical replicates. Each plot shows one random split. In some plots the points are equally distributed along the central line, but in other (bottom middle, right middle), it seems as if there are more hits on one group. The effect arises due to unequal split of the samples with more hits and the different number of hits comes from technical variation as all reactions were done with the same serum sample.

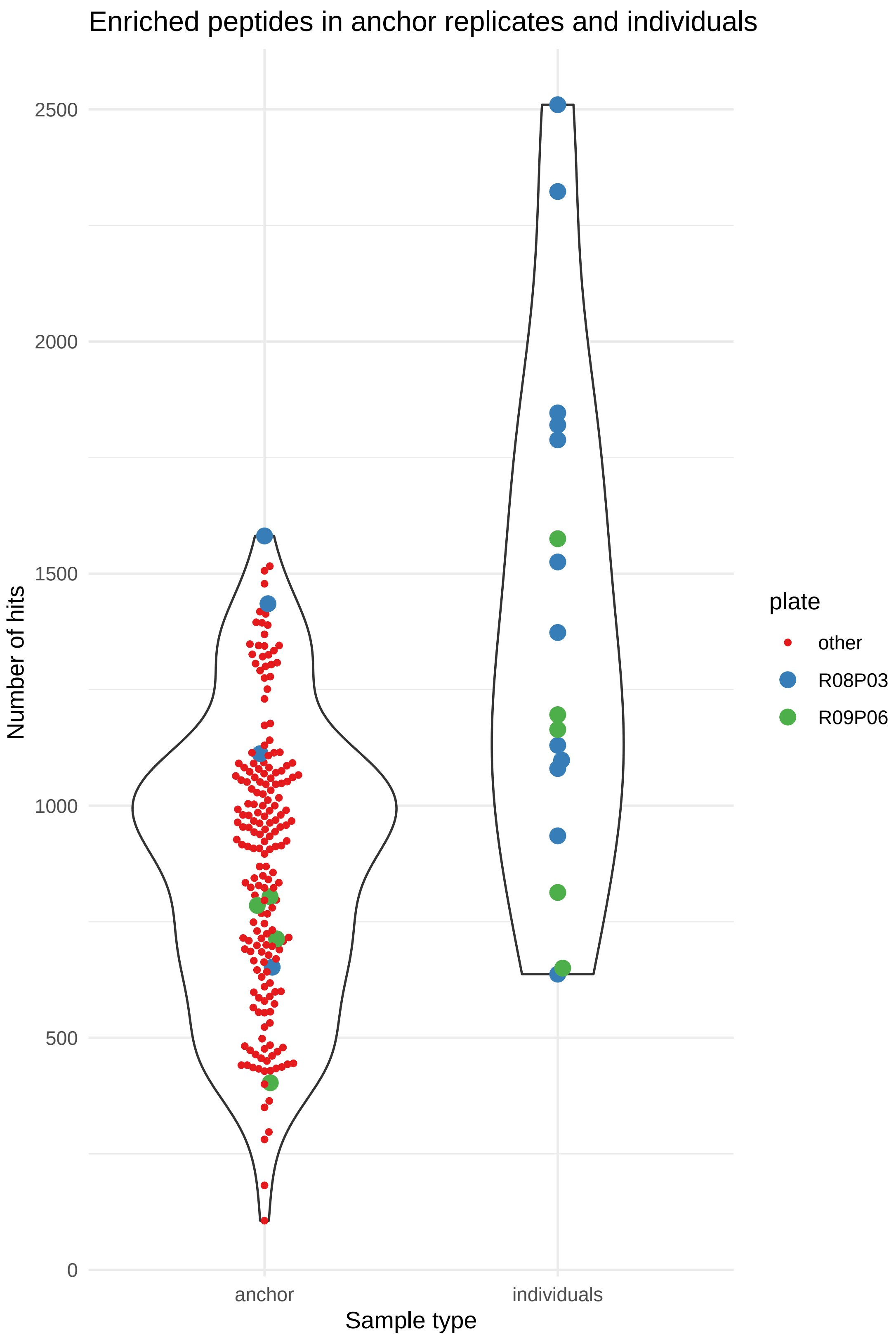


Supplementary Figure 6: Anchor sample replicates and the serum of the individuals that make the anchor serum have the comparable number of hits in the PhIP-Seq experiment, contrary to the expectation considering that a mixed serum contains more diverse antibody repertoire. The results suggest two things: 1) there is an upper bound of the number of hits our PhIP-Seq implementation can detect and 2) the composition and diversity of the antibodies in the serum influence set of retrieved bindings.

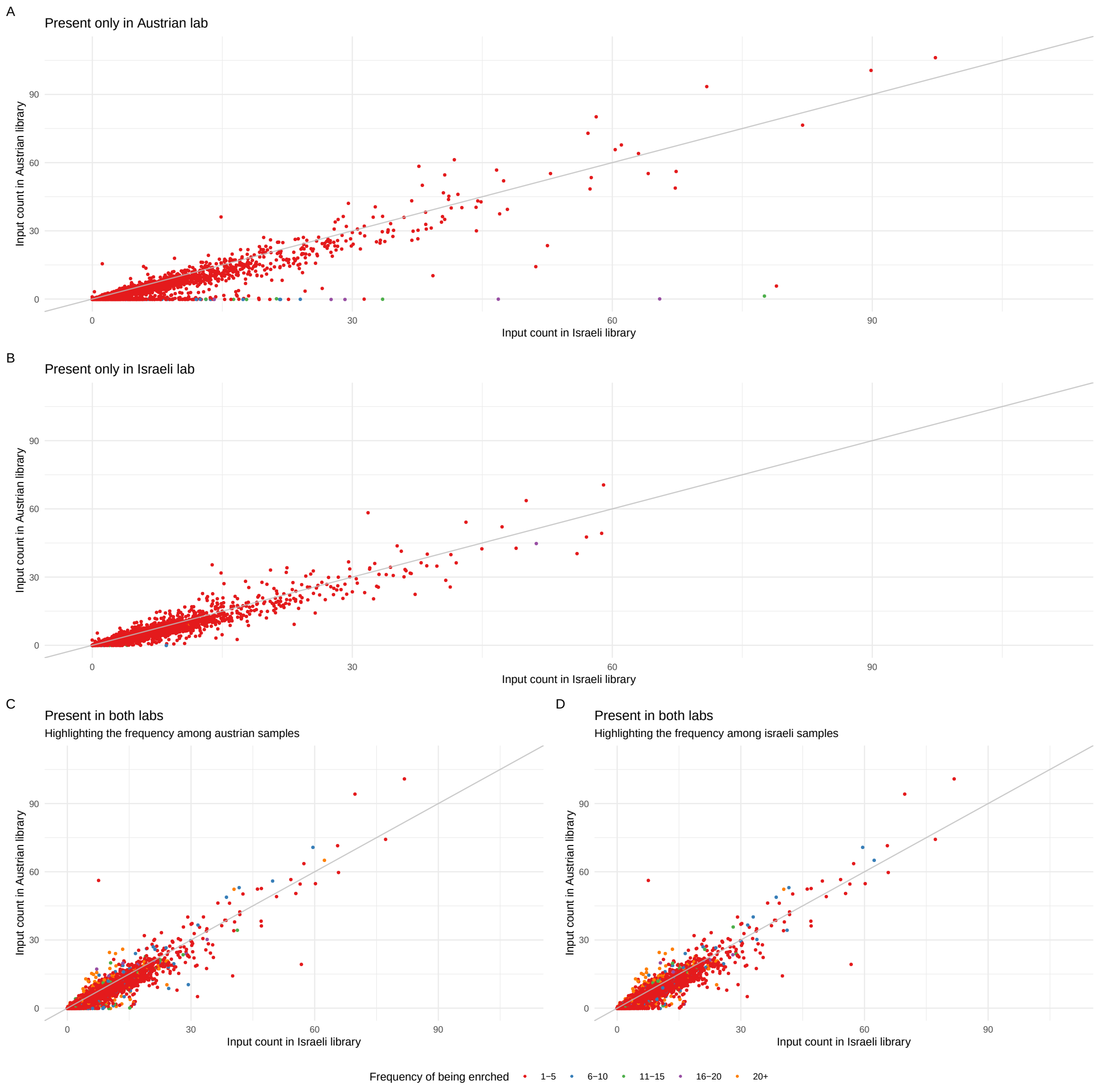


Supplementary Figure 7: Differences in the clone input counts in the Austrian and Israeli lab. Most of the clones show similar relative abundance, but some of the clones are detectable only in one or the other lab. **A:** Most of the clones have similar input counts at both labs, but some of them close to the x axis have much lower counts in Austrian lab. Regardless of the low counts, they are readily detected as enriched in samples. **B:** Among peptides detected as enriched only in Israeli lab, there is no obvious group of peptides with substantial difference in input counts. Moreover, almost all peptides that are exclusive for the Israeli lab are only rarely detected as enriched among samples (5 or less times). **C and D:** Input counts for peptides detected in both labs, but by colors showing the frequency of enrichment in one or the other lab. There is no obvious relationship between the frequency of enrichment in one or the other lab and the input counts that would point to the systematic differences. Given the results above, difference in the input counts doesn't seem to be the main driver of the differential peptide enrichment between two labs.
